# Effects of age in the strategic control of recollected content as reflected by modulation of scene reinstatement

**DOI:** 10.1101/2025.01.16.633416

**Authors:** Marianne de Chastelaine, Sabina Srokova, Sarah Monier, Joshua M. Olivier, Michael D. Rugg

**Affiliations:** Center for Vital Longevity and School of Behavioral and Brain Sciences, The University of Texas at Dallas, Dallas, TX 75235, USA; Department of Psychology and Evelyn F. McKnight Brain Institute, University of Arizona, Tucson, AZ 85721

**Keywords:** aging, cortical reinstatement, episodic memory, fMRI, recollection, retrieval gating

## Abstract

A previous study employing fMRI measures of retrieval-related cortical reinstatement reported that young, but not older, adults employ ‘retrieval gating’ to attenuate aspects of an episodic memory that are irrelevant to the retrieval goal. We examined whether the weak memories of the older adults in that study rendered goal-irrelevant memories insufficiently intrusive to motivate retrieval gating. Young and older participants studied words superimposed on rural or urban scenes, or on pixelated backgrounds. To strengthen memory for background information, word-image pairs were studied twice, initially centrally, and then in one of three locations. During scanning, one retrieval test probed memory for the test words’ studied backgrounds and another test assessed memory for their location. Background memory performance was markedly higher than in the prior study. Retrieval gating was examined in two scene-selective regions of interest, the parahippocampal place area (PPA) and the medial place area (MPA). In the background task, robust retrieval-related scene reinstatement effects were identified in both age groups. These effects were attenuated (‘gated’) in the location task in the young age group only, replicating the prior finding. The results did not differ when the two groups were sub-sampled to match strength of scene reinstatement when scene information was goal relevant. The findings indicate that older adults’ failure to gate goal-irrelevant scene information does not reflect age differences in memory strength and may instead reflect an age-related decline in top-down inhibitory control.

## 1. Introduction

Neuroimaging studies have often implicated encoding processes in age-related episodic memory decline (see Maillet and Rajah, 2014, for a meta-analysis; Wang and Cabeza, 2016, for review), whereas many of the neural correlates of retrieval processing appear to be relatively stable throughout much of the adult lifespan (e.g., Dulas and Duarte, 2011; de Chastelaine et al., 2016; de Chastelaine et al., 2017). For example, fMRI studies have reported retrieval effects in the ‘core recollection network’ (Kim, 2010; Rugg & Vilberg, 2013; King et al., 2015) to be of equal or near equal magnitude in young and older participants (e.g., Duarte at al., 2008; Dulas & Duarte, 2012; 2016; Wang & Giovanello, 2016; Wang et al., 2016; Hou et al., 2021). Furthermore, the neural correlates of retrieval processes thought to place high demands on executive control, such as post-retrieval monitoring, also appear to be minimally affected in healthy aging (e.g., Fandakova et al, 2013; Dulas and Duarte, 2014; Wang et al., 2016; Horne et al., 2021). Older adults’ adoption of specific ‘retrieval orientations’ – task sets that bias retrieval cue processing in service of a retrieval goal (Rugg, 2004) – appears, however, to be less spontaneous than is the case in young adults and may occur only when the retrieval task explicitly requires recollection of episodic content (Morcom and Rugg, 2004; Jacoby et al., 2005; Duverne et al., 2009).

Findings from a recent fMRI study investigating episodic memory retrieval in young and older adults suggest that another aspect of retrieval processing, ‘retrieval gating’, might be particularly vulnerable to the effects of increasing age (Srokova et al., 2021). Retrieval gating refers to the ability to control the contents of a retrieved memory so as to align them with the retrieval goal (Elward and Rugg, 2015; Elward et al., 2021). Retrieval gating has been proposed to depend on a combination of attentional suppression and amplification, whereby gating weakens cortical representations of goal-irrelevant information, especially when the information might interfere or compete with goal-relevant information, while strengthening the representation of relevant information (Elward and Rugg, 2015). In these prior studies, and in the present study also, the amount of retrieved content was indexed by the strength of retrieval-related cortical reinstatement, that is, the strength of reactivation of content-selective patterns of neural activity that were elicited during the encoding of a study event (for review, see Rugg and Srokova, 2024). On the widely held assumption that reinstatement supplies the content of a retrieved memory (Rugg and Srokova, 2024), retrieval gating can be operationalized in terms of the goal-dependent modulation of reinstatement.

In the study of Srokova et al. (2021), retrieval gating was examined in samples of young and older adults using an experimental procedure closely similar to that employed by Elward and colleagues (Elward and Rugg 2015; Elward et al., 2021). Words were presented at study superimposed over an image of an urban scene, a rural scene or a scrambled background. The word-image pairs appeared at one of three locations on the presentation monitor. The subsequent scanned memory test comprised two sub-tasks which were administered in short interleaved ‘mini-blocks’. In the ‘location’ task, the requirement was to judge the location in which recognized test words had been studied. In the ‘background’ task, the source memory judgment was with respect to the nature of the image that had been paired with the test word. In both age groups, retrieval-related scene reinstatement (i.e., greater neural activity for words studied with scenes than with a scrambled background) was identified in the background task in two scene-selective cortical regions, the parahippocampal place area (PPA) and medial place area (MPA; also referred to as the retrosplenial complex). Replicating prior findings (Elward and Rugg, 2015; Elward et al., 2021), reinstatement effects in the young participants were markedly attenuated in the location task - the hallmark of retrieval gating. Crucially, this finding was not evident in the older adults in whom reinstatement effects did not significantly differ between the two memory tasks. Additionally, in the young age group only, PPA scene reinstatement in the background task correlated positively across participants with source memory performance in that task, as well as with item memory more generally.

One interpretation of the absent retrieval gating in older adults reported by Srokova et al. (2021) is that the ability to control the content of recollected information declines with age. This interpretation is consistent with other evidence pointing to an age-related decline in cognitive control (e.g., Yuan and Raz, 2014; Zanto and Gazzaley, 2019), including the kinds of inhibitory control processes that might play a role in ‘suppressing’ otherwise memorable mnemonic content (Gazzaley et al., 2005, 2008; Chadick et al., 2014; Campbell et al., 2020; Weeks et al., 2020). It is noteworthy, however, that Srokova et al. (2021) reported that source memory accuracy was markedly lower in the older than the young age group. The relatively weak contextual memory in the older participants raises the possibility that their failure to demonstrate retrieval gating in the location task (i.e., when the background images were goal-irrelevant) did not reflect an inability to engage a gating strategy but, rather, a lack of motivation to do so. By this argument, the relatively weak memories for the background images were insufficiently intrusive to motivate engagement of retrieval gating when background information was goal irrelevant. This possibility is reminiscent of the finding that the adoption of specific retrieval orientations in older adults occurs only when there is a motivation to do so (Duverne et al., 2009). We examined this possibility in the present study by employing the same experimental design as Srokova et al. (2021), but with a study manipulation that strengthened memory for the background contexts so that performance in the older adults matched that of the younger adults in the prior study. If the failure in that study to identify retrieval gating effects in the older age group does indeed reflect weak memory for the background images, then we would expect to find evidence for the engagement of a gating strategy in the older age group employed in the present study.

As in Srokova et al. (2021), young and older adults studied a series of concrete nouns superimposed on a rural scene, an urban scene, or a scrambled background. Each word-image pair was studied twice, in contrast with the single study presentations employed previously. The subsequent scanned test phase was identical to that employed in the prior study: two interleaved memory tasks were employed, one of which probed memory for the test word’s studied background while the other tested location memory. We expected to identify scene reinstatement effects in the PPA and MPA in both age groups. We also expected that younger adults would demonstrate a retrieval gating effect in the form of attenuated scene reinstatement in the location task relative to the background task. As noted previously, if the failure of older adults to demonstrate retrieval gating in the study of Srokova et al. (2021) reflected a lack of incentive to employ gating because of weak memory for the backgrounds, the strengthening of these memories should motivate the adoption of a gating strategy and, hence, lead to attenuated scene reinstatement in the location task.

## 2. Materials and methods

### 2.1. Participants

Twenty-four young (18-31 years) and 24 older (63-79 years) cognitively healthy adults participated in the experiment. The participants were recruited from The University of Texas at Dallas and its surrounding communities. They were right-handed, had normal or corrected-to-normal vision, and were fluent English speakers from childhood. Participants had no history of neurological or psychiatric disease, substance abuse, diabetes, or recent use of prescription medication affecting the central nervous system. Participants were accepted into the study according to a set of inclusion and exclusion criteria intended to minimize the likelihood of including individuals with cognitive impairment arising from neuropathology, as determined by the neuropsychological test battery (see Neuropsychological testing, section 2.2, below). Data collected from 3 additional individuals (1 young and 2 older) were excluded because of poor performance on the experimental task (i.e., if the probability of source recollection – see section 2.5.2. – was less than 0.1). Participants gave informed consent in accordance with the UT Dallas Institutional Review Board and were compensated at the rate of $30 per hour for their time and $0.50 per mile for travel.

### 2.2. Neuropsychological testing

Prior to the experimental MRI session, participants completed a neuropsychological test battery which assessed a range of cognitive functions known either to decline or to be maintained with age. The Mini-Mental State Examination (MMSE) was employed to screen participants for dementia using a cutoff score of 26/30. The battery also comprised the California Verbal Learning Test-II (CVLT; Delis et al., 2000), Wechsler Logical Memory Tests 1 and 2 (Wechsler, 2009), the Forward and Backward Digit Span subtests of the Wechsler Adult Intelligence Scale Revised (WAIS-R; Wechsler, 1981), the Symbol Digit Modalities Test (SDMT; Smith, 1982), Trail Making Tests A and B (Reitan and Wolfson, 1985), the F-A-S subtest of the Neurosensory Center Comprehensive Evaluation for Aphasia (NCCEA; Spreen and Benton, 1977), the Category Fluency Test (Benton, 1968) the Test of Premorbid Functioning (TOPF; Wechsler, 2011) and Raven’s Progressive Matrices (short version; Raven et al., 2000). Participants were excluded prior to the MRI session if they performed > 1.5 SDs below the age norm on at least one memory-based test or on at least two non-memory tests, or if they obtained a standard score of < 100 as indexed by performance on the TOPF.

Visual acuity was tested with corrective lenses (if prescribed) using the Early Treatment Diabetic Retinopathy Study (ETDRS) charts and quantified with the logMAR metric (Ferris et al., 1982; Bailey and Lovie-Kitchin, 2013). All behavioral and fMRI analyses reported below were run both with and without visual acuity entered as a covariate. We report the results that did not include visual acuity as a covariate and note whenever its inclusion altered the results.

### 2.3. Materials

Critical stimuli comprised a pool of 240 visually presented words and 180 colored images depicting 60 urban and 60 rural scenes and 60 scrambled backgrounds. Scrambled images were created by randomly shuffling the pixels of 30 rural and 30 urban scenes. The words were high frequency concrete nouns selected from the word association norms compiled by Nelson et al. (2004). Study items included 180 of these words, superimposed over one of the scenes or scrambled backgrounds. All 180 study items were presented during an initial study phase (study phase 1) and were then re-presented, in a different order, during a second study phase (study phase 2). Whereas all study phase 1 items were presented in the center of the screen, only one third of the study phase 2 items were located centrally, with another third located on the left, and another third on the right of the screen. Test items included the 180 studied words (‘old’ items) intermixed with the 60 unstudied words remaining from the initial pool (‘new’ items). For each study-test stimulus list set, words were randomized such that all words from the pool were equally likely to be assigned to each background and location context at study or as a new item at test. For each study and test list, the different categories of items were pseudo-randomized so that no more than three items from the same category occurred successively. One buffer item was placed at the start of each study and test block. In total, 24 study-test stimulus list sets were created, one set per yoked pair of participants (one young and one older participant). Practice study and test lists were formed from items additional to the critical stimuli.

### 2.4. Experimental procedure

Figures 1 and 2 provide schematic depictions of the study and test tasks, respectively. Prior to scanning, participants were given instructions and a practice session for the study task and then completed the two experimental study phases on a laptop computer. A 5-minute rest break was provided between the two study phases. Each study phase included the sequential presentation of the 180 word-image pairs. These were divided into three study blocks separated by brief rest intervals. The study task (Figure 1) required participants to imagine a scenario in which the object denoted by the word moved around in or interacted with the background. Instructions were to press one of three buttons on a button box to indicate the vividness of the resulting mental image (‘1 = not vivid – 2 = somewhat vivid – 3 = very vivid’) using, respectively, the index, middle and ring fingers of their right hand. The rating cues were presented at the bottom of the display monitor along with a task reminder, ‘Imagine the object interacting with the background image’. During study phase 1, the word-image pairs were presented in the center of the screen. During study phase 2, the same word-image pairs were re-presented in a different presentation order. In contrast to study phase 1, the pairs were presented in one of three possible locations, namely on the left or right of the screen or in the center of the screen. For each study phase 2 stimulus list, equal numbers of images (20) from each image category (rural, urban and scrambled) were randomly assigned to each of the three screen locations (left, center or right).

**Figure 1.**
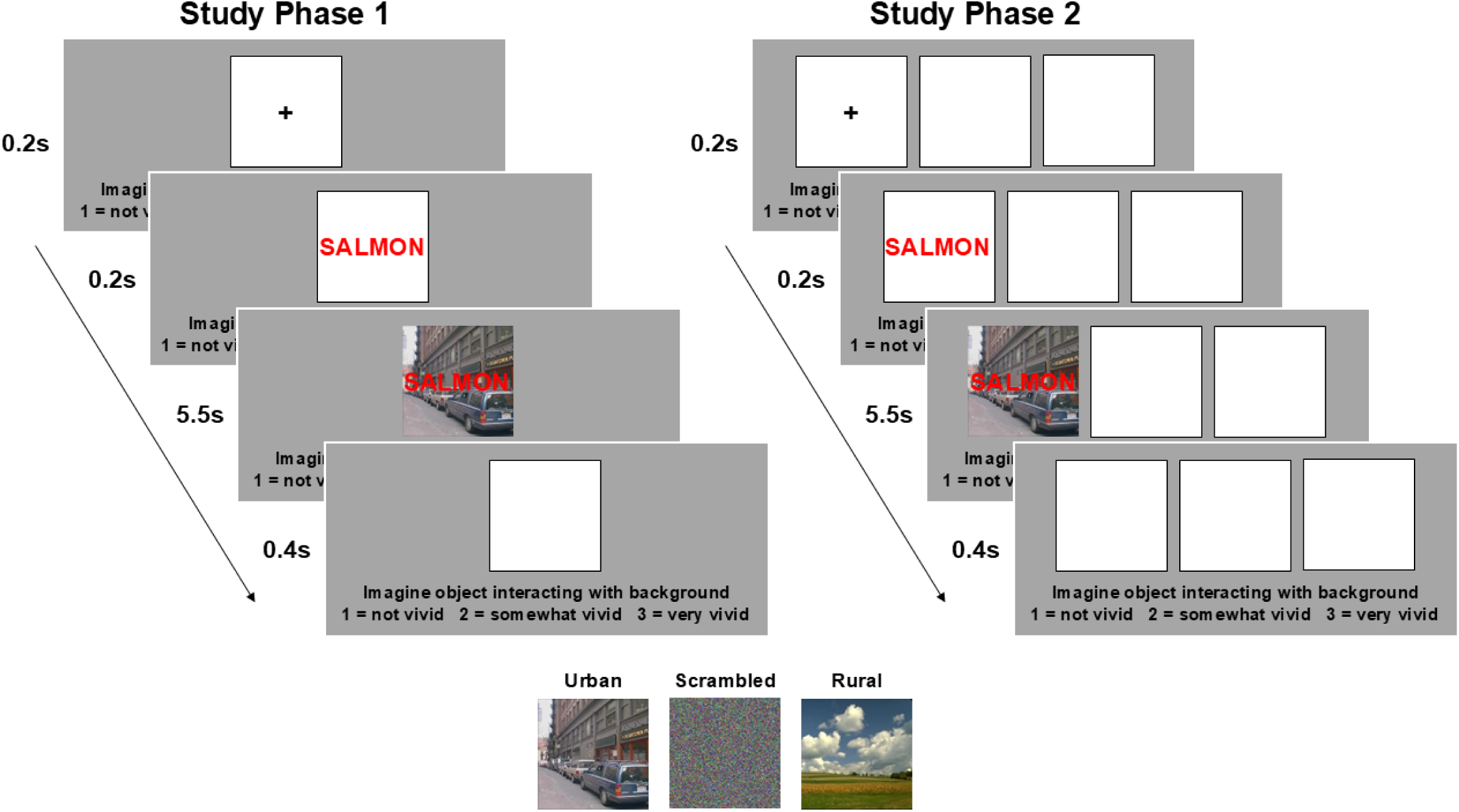
Schematic depiction of the two unscanned study phases. Timings denote the duration of each event within a trial.

**Figure 2.**
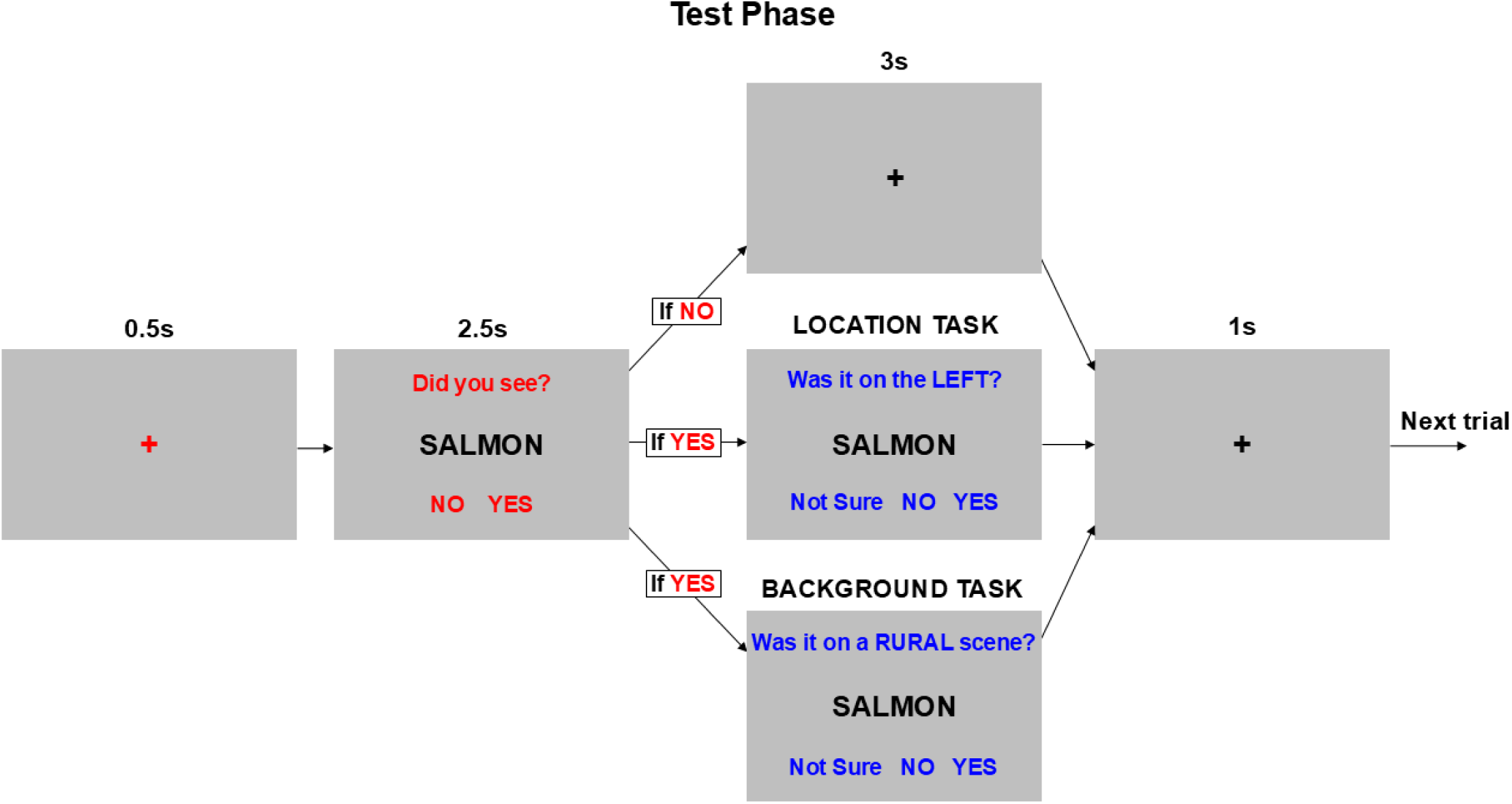
Schematic depiction of scanned test phase. Timings denote the duration of each event within a trial. The test phase included an initial recognition (item memory) question (red font). If participants indicated that they recognized the word, they were cued to answer an additional source memory question (blue font). For the location task, half of the participants were asked ‘Was it on the LEFT’ whereas the other half were asked ‘Was it on the RIGHT’. Similarly, for the background task, half of the participants were asked ‘Was it on a RURAL scene’ whereas the other half were asked ‘Was it on an URBAN scene’.

After completing study phase 2, participants were given instructions and a practice session for the test phase before entering the scanner. The test phase proper was completed inside the MRI scanner, starting around 40 minutes after completion of study phase 2. Test items were projected onto a translucent screen placed at the back of the scanner bore and viewed via a mirror mounted on the head coil. The test phase included sequential presentation of the 180 studied and 60 unstudied words organized into ten consecutive blocks – there were two blocks per scan run and, therefore, 5 scan runs in total (each scan run lasted 7.9 minutes). Equal numbers of test items from each contextual category (studied items paired with urban, rural and scrambled backgrounds, and at left, right and center locations) along with unstudied items were randomly assigned to each of the 10 test blocks. Each of the 5 scan runs, which were separated by brief rest intervals, included 1 block each of the two source memory tasks (i.e., the location and background tasks). Task order was constant across scan runs for each participant but counterbalanced across yoked pairs of young and older participants.

Prior to the start of each task block, participants were reminded with a visual cue which source memory task they were about to complete. For both tasks, the test words were initially presented with the memory prompt ‘Did you see?’ positioned above the word and the response cues ‘YES – NO’ below the word. Participants indicated whether or not they saw the word at study using either the index and middle fingers or the middle and ring fingers of the right hand. They were instructed to make a ‘YES’ response only when they were confident that they had seen the word at study and, if unconfident, to respond ‘NO’. When a word was identified as unstudied (i.e., a ‘NO’ response), a delay ensued until the next test word was presented. If a word was identified as studied (i.e., a ‘YES’ response), the initial prompt and response cues were replaced with a prompt and response cues pertaining to the appropriate source memory question about either the location or background contexts of the studied word (see Figure 2).

For the location task, participants were required to indicate the location of the word when it was presented in the second study block in response to one of two possible prompts that specified the ‘target’ location presented above the word. For half of the participants, the prompt asked ‘Was it on the LEFT?’ while, for the remainder of the participants, the prompt asked ‘Was it on the RIGHT?’. For the background task, the requirement was to indicate which class of background image the word was studied with in response to one of two possible prompts that specified the ‘target’ background. For half of the participants, the prompt asked ‘Was it on an URBAN scene?’ while, for the remainder of the participants, the prompt asked ‘Was it on a RURAL scene?’. Target location (left or right) and target scene (urban or rural) contexts remained constant across scan runs for each participant. The center location for the location task and the scrambled image for the background task were never employed as prompts and, therefore, were always non-target contexts. For both tasks, the response cue ‘YES – NO – Not Sure’ was presented below the word. Thus, items that required a ‘YES’ source response were designated as ‘target’ items whereas those that required a ‘NO’ source response were designated as ‘non-target’ items. Participants were instructed to respond ‘YES’ and ‘NO’ only when they were confident that the word had or had not been paired, respectively, with the target context at study and, if unconfident, to respond ‘Not Sure’. Participants indicated their response with the index, middle and ring fingers of their right hand. Response-finger mapping was counterbalanced across participants with the constraint that the ‘Not Sure’ response was mapped onto either the index or ring finger. For each participant, the fingers used for the ‘YES’ and ‘NO’ responses were the same for both the item memory (‘Did you see?’) and follow-up source memory prompts.

Each study trial began with the presentation of a black fixation cross in the center of a black-bordered white square, displayed on a grey background for 200 ms at the location assigned to the upcoming word-image pair. For study phase 1, a single square was displayed in the center of the screen whereas, for study phase 2, three identical squares were displayed, one to the left and one to the right of the screen and one in the center of the screen. The squares remained on the screen throughout each trial. The study word, presented in red Arial font, replaced the cross for 200 ms to ensure participants could clearly see the word prior to being superimposed over its paired image. The word-image pairs were then displayed for 5500 ms during which time the vividness rating was required. The study pairs were then removed from the screen leaving the square(s) on screen for another 400 ms. The response cues and study task reminder were displayed in black font at the bottom of the screen throughout the trial.

Each test trial began with the presentation of a red fixation cross in the center of a grey screen for 500 ms. The cross was then replaced by the test word, displayed in black Arial font, along with the initial item memory prompt (‘Did you see?’) and cues (‘YES – NO’) shown in red font, for 2500 ms. If participants indicated they did not recognize the word from the study phase (i.e., a ‘NO’ response), the word, prompts and cues disappeared and a black fixation cross was displayed in the center of the screen for 4000 ms. When participants indicated that they did recognize the word from study (i.e., a ‘YES’ response), the word remained on the screen, along with the source memory prompt and response cues in blue font, for 3000 ms. The word, prompts and cues were then replaced with a black fixation cross in the center of the screen for 1000 ms. Test items were intermixed with 80 null trials (8 per test block with no more than 2 null trials in succession) that comprised the presentation of a black fixation cross against the grey screen for 6500 ms.

For both the study and test tasks, participants were instructed to respond as quickly as possible without sacrificing accuracy. PsychoPy v2022.2.4 (Peirce et al., 2019) was used for stimulus presentation and recording participant responses.

### 2.5. Behavioral measures and analyses

#### 2.5.1. Study

Vividness ratings (1, not vivid, 2, somewhat vivid and 3, very vivid) and their associated RTs were averaged across study items of the same background context (target scenes, non-target scenes and scrambled images) for each study phase and participant. To determine if these measures differed according to age group, study phase or background context, the vividness ratings and RTs were each analyzed with a 2 (age group) x 2 (study phase) x 3 (background context) mixed effects ANOVA.

#### 2.5.2. Test

For both the location and background tasks, we computed estimates of item recognition accuracy (Pr) and source memory accuracy (pSR – i.e., probability of source recollection) for each participant. Pr was computed as the difference between the hit rate for studied words and the false alarm rate for unstudied words. pSR was computed using a modified single high-threshold model (Snodgrass and Corwin, 1988) that accounts for the guessing rate (e.g., Mattson et al., 2014). Thus, pSR was computed as:

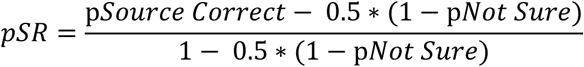

pSource Correct and pNot Sure variables refer to the proportion of hits that were accompanied by an accurate or ‘Not Sure’ source memory judgment, respectively.

Pr and pSR were each analyzed with a 2 (age group) x 2 (task) mixed effects ANOVA to identify any differences in these measures according to age group or source memory task (location versus background). Test RTs were averaged across item hits belonging to the same background context (target scenes, non-target scenes and scrambled images) separately for each task (location and background) and participant. The RTs were analyzed with a 2 (age group) x 2 (task) x 3 (background) mixed effects ANOVA.

As the aim of the current experiment was to boost source memory performance in the background task relative to that reported in Srokova et al. (2021), we also ran a between-experiments contrast of the pSR metrics from each experiment, using a 2 (experiment) x 2 (age group) x 2 (task) mixed effects ANOVA.

### 2.6. fMRI acquisition

A Siemens Prisma 3T MRI scanner equipped with a 32-channel head coil was used to acquire functional and anatomical images. A 3D MP-RAGE pulse sequence (TR = 2300 ms, TE = 2.41 ms, FOV = 256×256 mm, voxel size 1×1×1 mm, 160 slices, sagittal acquisition) was employed for T1-weighted anatomical image acquisition. Functional scans were acquired with a T2*–weighted echo-planar imaging sequence with a multiband factor of 3 and an acceleration factor of 2 (TR = 1.52 s, TE = 30 ms, voxel size = 2 × 2 × 2 mm, 0.4 mm inter-slice gap, 66 slices, anterior-to-posterior phase encoding direction, flip angle = 70°). A double-echo fieldmap sequence (TE 1 = 4.92 ms, TE 2 = 7.38 ms, TR = 652 ms, FOV 220 mm, 66 slices, 2 mm slice thickness, flip angle = 60°,) was acquired immediately after the last scan run of the test phase.

### 2.7. fMRI preprocessing

fMRI data were preprocessed and analyzed using SPM12 (http://www.fil.ion.ucl.ac.uk) and custom Matlab code (MathWorks). Functional images were field-map corrected, realigned, reoriented to the AC-PC line and spatially normalized using a sample-specific template. The template was created by first normalizing (Ashburner and Friston, 1999) the mean volume of each participant’s functional time series with reference to a standard EPI template based on the Montreal Neurological Institute (MNI) reference brain (Cocosco et al., 1997). The normalized images were averaged across participants to generate a template that was equally weighted with respect to the 2 age groups (de Chastelaine et al. 2011, 2016). The normalized images were then smoothed using a 5 mm full-width half-maximum (FWHM) Gaussian kernel. The functional data from the different test blocks were concatenated using the spm_concatenate.m function before estimating the first level GLMs.

### 2.8. ROIs

Figure 3 shows the four scene-selective ROIs employed in the current experiment (bilateral PPA and MPA). These ROIs were the same as those employed by Srokova et al. (2021). They were derived from the localizer task that was employed in that study and were the outcome of the conjunction between scenes > objects and scenes > scrambled images contrasts conducted at the group level. The contrasts took the form of simple effects of item type to ensure the ROIs were unbiased with respect to age group. The contrasts were masked anatomically to restrict the ROIs to parahippocampal/fusiform (PPA) and retrosplenial/medial occipital (MPA) cortex, as described in Srokova et al. (2021).

**Figure 3.**
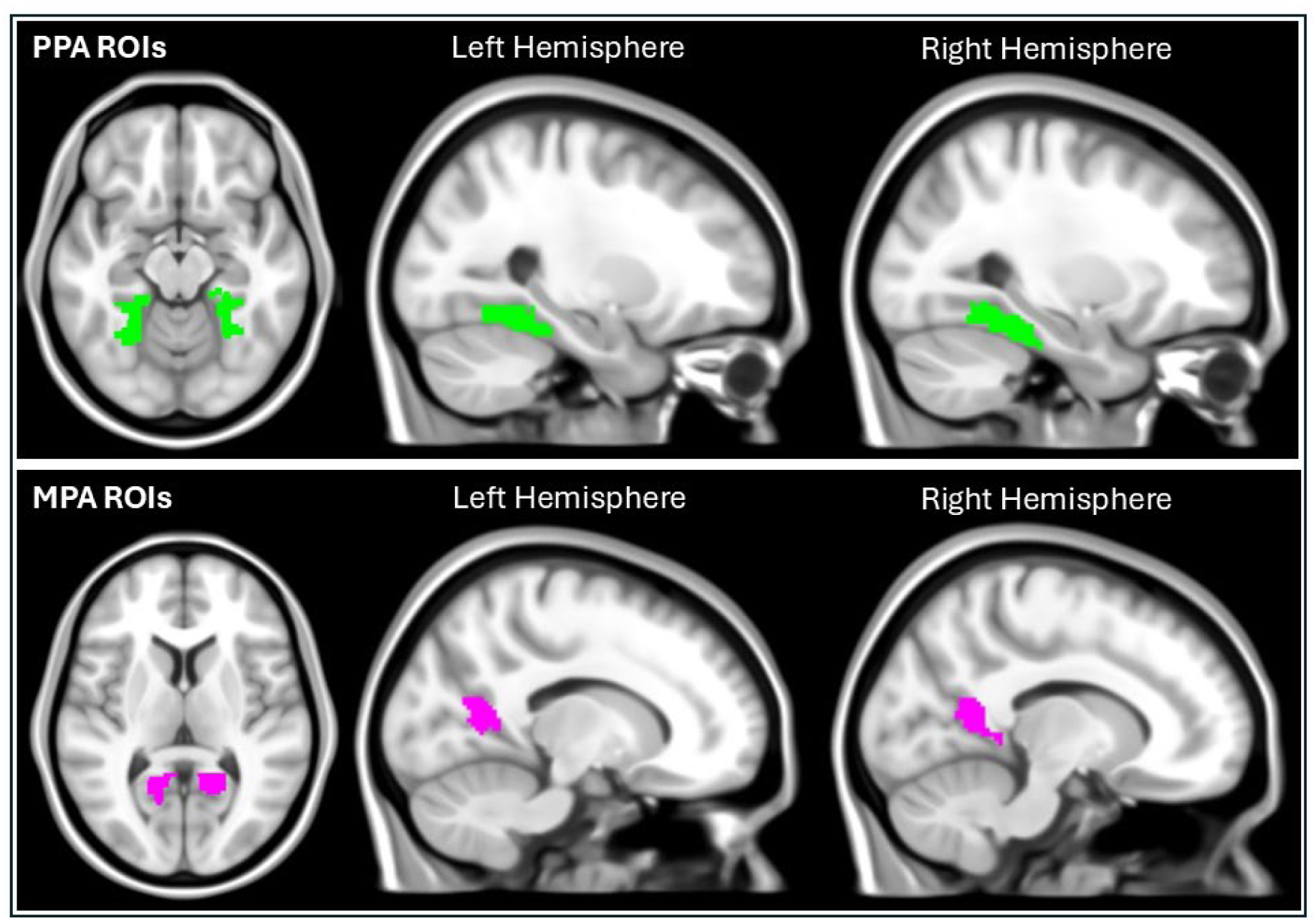
Bilateral scene-selective ROIs shown on a T1-weighted ICBM 152 MNI brain: PPA (upper panel); MPA (lower panel).

### 2.9. Reinstatement index

Single-trial parameter estimates of the magnitudes of item-elicited BOLD responses (beta estimates) were derived for each participant from a first level GLM implementing a ‘least-squares-all’ approach (Mumford et al., 2012; Abdulrahman and Henson, 2016). Neural activity elicited by each test item was modeled as a separate delta function convolved with a canonical HRF – thus, each test item was modeled as a separate event of interest. Six motion regressors and the four constants modeling the mean BOLD signal for scan runs 2-5 were included as covariates of no interest. As in Srokova et al. (2021) mean beta estimates from the bilateral PPA and MPA ROIs were used to compute *scene reinstatement indices* separately for the location and background tasks. The reinstatement index was estimated as the difference in mean BOLD amplitude for test words studied with scenes and those studied with scrambled backgrounds after scaling the difference by the pooled inter-item standard deviation:

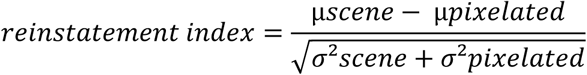

As is evident from the formula, an index greater than zero reflects greater mean BOLD amplitude for words studied with scenes than for words studied with scrambled backgrounds, indicative of cortical reinstatement of scene information. Importantly, because of the scaling function, the reinstatement index is insensitive to individual differences (and, by extension, age group differences) in the gain of the hemodynamic response function (HRF) that mediates the relationship between neural and BOLD activity (cf. Liu et al., 2013). As described previously (Elward and Rugg, 2015; Elward et al., 2021; Srokova et al., 2021), test items included in the computation of the reinstatement index comprised all items that were correctly recognized as having been studied (item hits) and that were associated with the ‘non-target’ image categories (i.e., either rural or urban scenes, depending upon the assigned target category, and scrambled images), regardless of source memory accuracy. Restricting the reinstatement metric to non-target backgrounds reduces the possible confounding effects of the multiple responses associated with the scene images (a ‘YES’ response for targets but a ‘NO’ response for non-targets) as opposed to the single response for scrambled backgrounds (always ‘NO’). Of importance, on the assumption that, in the absence of gating, scene retrieval would be equally likely in the background and location tasks, the inclusion of all correctly recognized items also allows for an unbiased comparison of scene reinstatement between the location and background tasks.

### 2.10. fMRI analyses

To identify differences in scene reinstatement according to age group or task, the indices were subjected to a 2 (age group) x 2 (task) x 2 (hemisphere) x 2 (ROI) mixed effects ANOVA. We also went on to directly compare the scene reinstatement indices obtained in the current experiment with those originally reported by Srokova et al. (2021). To effect this comparison, we ran a between experiments analysis using a 2 (experiment) x 2 (age group) x 2 (task) x 2 (ROI) x 2 (hemisphere) mixed effects ANOVA. To obtain a clearer understanding of the scene reinstatement effects across experiments, we also conducted secondary analyses of the mean amplitudes of the BOLD signals elicited by the test words studied with non-target scenes and the scrambled backgrounds, using a 2 (experiment) x 2 (age group) x 2 (background context: scenes, scrambled) x 2 (task) x 2 (ROI) x 2 (hemisphere) mixed effects ANOVA. Additionally, we conducted multiple regression analyses to examine the relationships between scene reinstatement effects in the PPA for the background task and memory performance for items (Pr) and background contexts (pSR). Following Srokova et al. (2021), as Pr scores for the background and location tasks were highly correlated (r = 0.867, p < 0.001) and did not significantly differ (p = 0.253), these were averaged to produce a single item memory score for each participant. Again, following Srokova et al. (2021), the initial regression models included age group, the reinstatement index and the age group by reinstatement interaction terms as predictors of either item or source memory performance. A subsequent pair of models employed the combined datasets, and predicted memory performance using as predictors the same terms as previously, along with the additional terms of experiment and the experiment by reinstatement index interaction.

### 2.11. Statistical analysis and software packages

Statistical analyses were performed using SPSS 29.0. The Greenhouse-Geisser correction was applied to ANOVA contrasts when appropriate. As in Srokova et al. (2021), t-tests were two-tailed except for the one-sample t-tests run to determine whether reinstatement indices were significantly greater than zero. Significance levels for all tests were set at p < 0.05. Effect sizes are reported as partial-η^2^ for ANOVAs and as Cohen’s d for t-tests. Figures plotting the fMRI data were created using the package ggplot2 (Wickham, 2016) with R software (R Core Team, 2020).

## 3. Results

### 3.1. Neuropsychological data

Table 1 summarizes demographic and neuropsychological data for the two age groups. The data were analyzed using independent t-tests to compare the two groups. In brief, for CVLT recognition hits were higher in the young compared to the older adults and false alarms were higher in the older adults relative to the young. Older participants also demonstrated poorer performance than the young group on tests of speeded cognition (SDMT and Trail Making Tests A and B) and on the Category Fluency Test. The performance of older participants was lower than that of younger participants on a test of fluid intelligence (Raven’s Progressive Matrices), while performance on a test of crystallized intelligence (TOPF) was age-invariant. Older adults had significantly more years of education than younger adults.

**Table 1.** Demographic data and neuropsychological test performance [mean (SD)] for the young and older groups, and p values from the results of independent t-tests comparing performance between the two age groups.

|  | Young | Older | p-value |
| --- | --- | --- | --- |
| N (male/female) | 9/15 | 6/18 |  |
| Age | 23.29 (3.77) | 69.33 (4.40) |  |
| Years of Education | 15.79 (2.04) | 17.46 (2.28) | 0.011 * |
| MMSE | 29.13 (1.12) | 29.13 (1.23) | 1.000 |
| CVLT Short Delay - Free | 12.50 (2.60) | 11.33 (2.50) | 0.120 |
| CVLT Short Delay - Cued | 13.04 (2.42) | 12.75 (2.03) | 0.653 |
| CVLT Long Delay - Free | 12.96 (2.26) | 12.21 (2.55) | 0.286 |
| CVLT Long Delay - Cued | 13.46 (2.11) | 12.71 (2.49) | 0.266 |
| CVLT Recognition Hits | 15.58 (0.65) | 14.96 (1.04) | 0.017 * |
| CVLT False Alarms | 0.75 (0.99) | 1.58 (1.72) | 0.047 * |
| F-A-S | 46.29 (13.10) | 43.88 (9.68) | 0.471 |
| WMS Logical Memory I | 28.63 (5.84) | 27.83 (5.35) | 0.627 |
| WMS Logical Memory II | 26.67 (6.76) | 25.25 (5.44) | 0.428 |
| Symbol Digit Modalities Test | 61.04 (9.22) | 48.25 (8.13) | < 0.001 * |
| Trails A (s) | 21.89 (5.15) | 28.00 (7.73) | 0.003 * |
| Trails B (s) | 48.06 (18.22) | 71.23 (27.46) | 0.001 * |
| Forward/Backward Digit Span | 18.17 (4.32) | 18.58 (4.14) | 0.735 |
| Category Fluency Test | 23.92 (5.17) | 20.79 (4.63) | 0.033 * |
| TOPF (Standard Score) | 117.46 (9.22) | 115.04 (8.23) | 0.421 |
| TOPF (Raw Score) | 55.33 (8.28) | 57.21 (7.71) | 0.343 |
| Raven's Progressive Matrices I | 10.54 (1.84) | 8.75 (2.09) | 0.003 * |

\*Significant difference at $p < 0.05$ .

### 3.2. Behavioral data

#### 3.2.1. Study phase

Table 2 shows group mean vividness ratings and RTs for study phases 1 and 2 categorized according to the background context (target scenes, nontarget scenes and scrambled background) of the study items. Vividness and RT data from block 3 of study phase 2 for one older adult were not recorded due to a technical malfunction – therefore, only data from blocks 1 and 2 from study phase 2 were included in the study phase 2 summary and analysis for that participant.

**Table 2.** Mean (SD) vividness ratings and RTs from the two study phases for the young and older groups.

|  | Young | Older |
| --- | --- | --- |
| ***Study Phase 1*** |  |  |
| Vividness Ratings |  |  |
| Target scene | 2.30 (0.43) | 2.49 (0.39) |
| Non-target scene | 2.30 (0.46) | 2.52 (0.34) |
| Scrambled background | 2.04 (0.51) | 2.32 (0.47) |
| RTs (ms) |  |  |
| Target scene | 3165 (816) | 3461 (711) |
| Non-target scene | 3146 (808) | 3435 (698) |
| Scrambled background | 3018 (843) | 3241 (695) |
| ***Study Phase 2*** |  |  |
| Vividness Ratings |  |  |
| Target scene | 2.38 (0.49) | 2.47 (0.33) |
| Non-target scene | 2.36 (0.50) | 2.42 (0.38) |
| Scrambled background | 2.07 (0.60) | 2.18 (0.57) |
| RTs (ms) |  |  |
| Target scene | 2750 (792) | 2929 (804) |
| Non-target scene | 2775 (832) | 2905 (812) |
| Scrambled background | 2668 (831) | 2840 (808) |

The study phase data were submitted to 2 (age group) x 2 (study phase) x 3 (background context) mixed effects ANOVAs. ANOVA of the vividness ratings revealed a main effect of background context [F(1.38,63.28) = 25.870, p < 0.001, partial η^2^ = 0.360]. Follow-up pairwise t-tests indicated that, across the 2 age groups and the 2 study phases, vividness ratings for words paired with scrambled backgrounds were lower than for words paired with either target scenes [t(47) = 5.409, p < 0.001, d = 0.781] or non-target scenes [t(47) = 5.453, p < 0.001, d = 0.787]. There was no significant difference in vividness ratings between target and non-target scenes (p = 0.565). The ANOVA also revealed a study phase by age group interaction [F(1,46) = 5.126, p = 0.028, partial η^2^ = 0.100], reflecting slight increases and decreases in mean vividness ratings from study phase 1 to study phase 2 in young and older adults, respectively. However, the differences in mean ratings between the 2 study phases were not statistically significant in either age group (young: p = 0.158; older: p = 0.096).

ANOVA of the RT data revealed a main effect of study phase [F(1,46) = 48.913, p < 0.001, partial η^2^ = 0.515] indicating, across background contexts and age groups, faster RTs in study phase 2 relative to study phase 1. There was also a main effect of background context [F(1.66,76.22) = 9.519, p < 0.001, partial η^2^ = 0.171] and a phase by background interaction [F(2,92) = 3.512, p = 0.034, partial η^2^ = 0.071]. ANOVAs conducted separately for the two study phases indicated a main effect of background context for study phase 1 items [F(1.65,75.86) = 18.531, p < 0.001, partial η^2^ = 0.287] but not for study phase 2 items (p = 0.1). Follow-up pairwise t-tests indicated that RTs were faster for study phase 1 items paired with scrambled backgrounds relative to those paired either with target [t(47) = 4.898, p < 0.001, d = 0.707] or non-target scenes [t(47) = 4.566, p < 0.001, d = 0.659]. There were no differences in RTs to items paired with target versus non-target scenes (p = 0.350).

#### 3.2.2. Test phase

Table 3 summarizes memory performance for the test phase (upper panel for the current experiment). Separate 2 (age group) x 2 (task) ANOVAs were conducted on the item memory (Pr) and source memory (pSR) metrics. For Pr, the ANOVA revealed a main effect of age group [F(1,46) = 4.239, p = 0.045, partial η^2^ = 0.084], indicating better item memory performance for the young relative to the older participants. However, the effect of age group was no longer significant when visual acuity was added as a covariate in the ANOVA model (p = 0.091). There were no significant effects or interactions involving the factor of task. For pSR, the ANOVA similarly revealed a main effect of age group [F(1,46) = 23.898, p < 0.001, partial η^2^ = 0.342], indicating better source memory performance for the young group relative to the older participants. There was also a main effect of task [F(1,46) = 250.364, p < 0.001, partial η^2^ = 0.845], indicating better source memory performance on the background task relative to the location task.

**Table 3.** Test performance [mean (SD)] for the location and background tasks for the young and older groups for the current study (upper panel) and from Srokova et al. (2021) (lower panel).

|  | Young |  | Older |  |
| --- | --- | --- | --- | --- |
| Current Experiment |  |  |  |  |
| Test Phase | Location | Background | Location | Background |
| Item Hit Rate | 0.90 (0.09) | 0.90 (0.10) | 0.82 (0.14) | 0.81 (0.14) |
| False Alarm Rate | 0.08 (0.08) | 0.09 (0.12) | 0.09 (0.13) | 0.10 (0.11) |
| Item Memory (Pr) | 0.82 (0.12) | 0.81 (0.16) | 0.73 (0.18) | 0.72 (0.16) |
| Source Memory (pSR) | 0.29 (0.29) | 0.74 (0.15) | 0.08 (0.13) | 0.53 (0.23) |
| Proportion Source Correct |  |  |  |  |
| Target scene | 0.51 (0.21) | 0.85 (0.11) | 0.34 (0.21) | 0.71 (0.16) |
| Non-target scene | 0.52 (0.18) | 0.84 (0.13) | 0.26 (0.19) | 0.77 (0.18) |
| Scrambled background | 0.50 (0.21) | 0.87 (0.09) | 0.26 (0.20) | 0.71 (0.20) |
| Proportion Not Sure |  |  |  |  |
| Target scene | 0.36 (0.20) | 0.07 (0.07) | 0.49 (0.27) | 0.09 (0.12) |
| Non-target scene | 0.32 (0.20) | 0.10 (0.11) | 0.55 (0.28) | 0.12 (0.18) |
| Scrambled background | 0.37 (0.20) | 0.09 (0.10) | 0.58 (0.29) | 0.13 (0.16) |
| Test RTs (ms) – Item Hits |  |  |  |  |
| Target scene | 1171 (105) | 1133 (133) | 1421 (157) | 1419 (166) |
| Non-target scene | 1189 (119) | 1148 (121) | 1411 (146) | 1440 (163) |
| Scrambled background | 1181 (119) | 1158 (122) | 1446 (178) | 1441 (148) |
| Srokova et al. (2021) |  |  |  |  |
| Test Phase | Location | Background | Location | Background |
| Item Hit Rate | 0.81 (0.12) | 0.80 (0.14) | 0.72 (0.12) | 0.71 (0.10) |
| False Alarm Rate | 0.15 (0.15) | 0.16 (0.12) | 0.14 (0.09) | 0.12 (0.08) |
| Item Memory (Pr) | 0.66 (0.21) | 0.64 (0.18) | 0.58 (0.15) | 0.59 (0.15) |
| Source Memory (pSR) | 0.23 (0.16) | 0.49 (0.19) | 0.16 (0.13) | 0.38 (0.19) |
| Proportion Source Correct |  |  |  |  |
| Target scene | 0.48 (0.15) | 0.70 (0.17) | 0.41 (0.15) | 0.65 (0.16) |
| Non-target scene | 0.49 (0.17) | 0.73 (0.17) | 0.39 (0.15) | 0.66 (0.19) |
| Scrambled background | 0.52 (0.17) | 0.63 (0.21) | 0.47 (0.18) | 0.56 (0.25) |
| Proportion Not Sure |  |  |  |  |
| Target scene | 0.28 (0.18) | 0.15 (0.12) | 0.33 (0.19) | 0.14 (0.13) |
| Non-target scene | 0.27 (0.17) | 0.16 (0.15) | 0.33 (0.22) | 0.14 (0.16) |
| Scrambled background | 0.31 (0.18) | 0.25 (0.18) | 0.33 (0.22) | 0.18 (0.18) |

We employed a 2 (age group) x 2 (task) x 3 (background) mixed effects ANOVA to examine RTs for item hits. The ANOVA revealed a main effect of age group [F(1,46) = 54.352, p < 0.001, partial η^2^ = 0.542], reflecting slower RTs for older relative to young adults. There were no other significant main effects or interactions.

##### 3.2.2.1. Between experiment analysis

As the aim of the current experiment was to boost source memory performance in the background task relative to that reported in Srokova et al. (2021), we ran a between-experiments analysis on the pSR measures to verify that we achieved that aim (see Table 3, lower panel, for a summary of the behavioral data from the prior study). A 2 (experiment) x 2 (age group) x 2 (task) ANOVA revealed a main effect of task [F(1,88) = 277.384, p < 0.001, partial η^2^ = 0.759] indicating, across experiments and age groups, better source memory performance for background relative to location information. There were also main effects of age group [F(1,88) = 26.332, p < 0.001, partial η^2^ = 0.230] and experiment [F(1,88) = 10.346, p = 0.002, partial η^2^ = 0.105] indicating, respectively, better source memory performance overall for the young relative to the older participants and for the current experiment relative to its predecessor. Crucially, the ANOVA also revealed an interaction between experiment and task [F(1,88) = 25.541, p < 0.001, partial η^2^ = 0.225]. Follow-up independent t-tests for each task indicated that the increase in source memory performance from the previous to the current experiment was confined to the background task [t(90) = 4.768, p < 0.001, d = 0.995; for location information, p = 0.879]. As can be seen in Table 3, the older participants in the current study demonstrated numerically higher pSR scores for the background task than the young participants from the previous study. Thus, we achieved our aim of boosting memory performance in older adults in the current study to equal the performance of the young adults in the previous study.

### 3.3. fMRI results

#### 3.3.1. Reinstatement index

Summary statistics for the reinstatement indices, segregated by age group, ROI and task, are given in Table 4 (upper panel) and illustrated in Figure 4A. The data were subjected to a 2 (age group) x 2 (task) x 2 (hemisphere) x 2 (ROI) mixed effects ANOVA. The ANOVA revealed a main effect of task [F(1,46) = 5.152, p = 0.028, partial η^2^ = 0.101], indicating greater scene reinstatement in the background task relative to the location task, and a main effect of ROI [F(1,46) = 8.473, p = 0.006, partial η^2^ = 0.156], reflecting greater reinstatement in the MPA than in the PPA. The age group by task interaction did not reach significance [F(1,46) = 2.965, p = 0.092, partial η^2^ = 0.061].

**Table 4.** Mean (SD) reinstatement estimates from the PPA and MPA ROIs for the location and background tasks according to age group and experiment.

|  | Young | Older |
| --- | --- | --- |
| ***Current Experiment*** |  |  |
| PPA location task | 0.0612 (0.1903) | 0.0999 (0.3343) |
| MPA location task | 0.1342 (0.2381) | 0.1523 (0.3236) |
| PPA background task | 0.2574 (0.3607) | 0.1469 (0.2317) |
| MPA background task | 0.3499 (0.2951) | 0.1620 (0.3027) |
| ***Srokova et al. (2021)*** |  |  |
| PPA location task | 0.1151 (0.2648) | 0.1603 (0.2170) |
| MPA location task | 0.1129 (0.2954) | 0.2751 (0.3156) |
| PPA background task | 0.3199 (0.3440) | 0.0983 (0.3714) |
| MPA background task | 0.3101 (0.1956) | 0.1579 (0.3506) |

**Figure 4.**
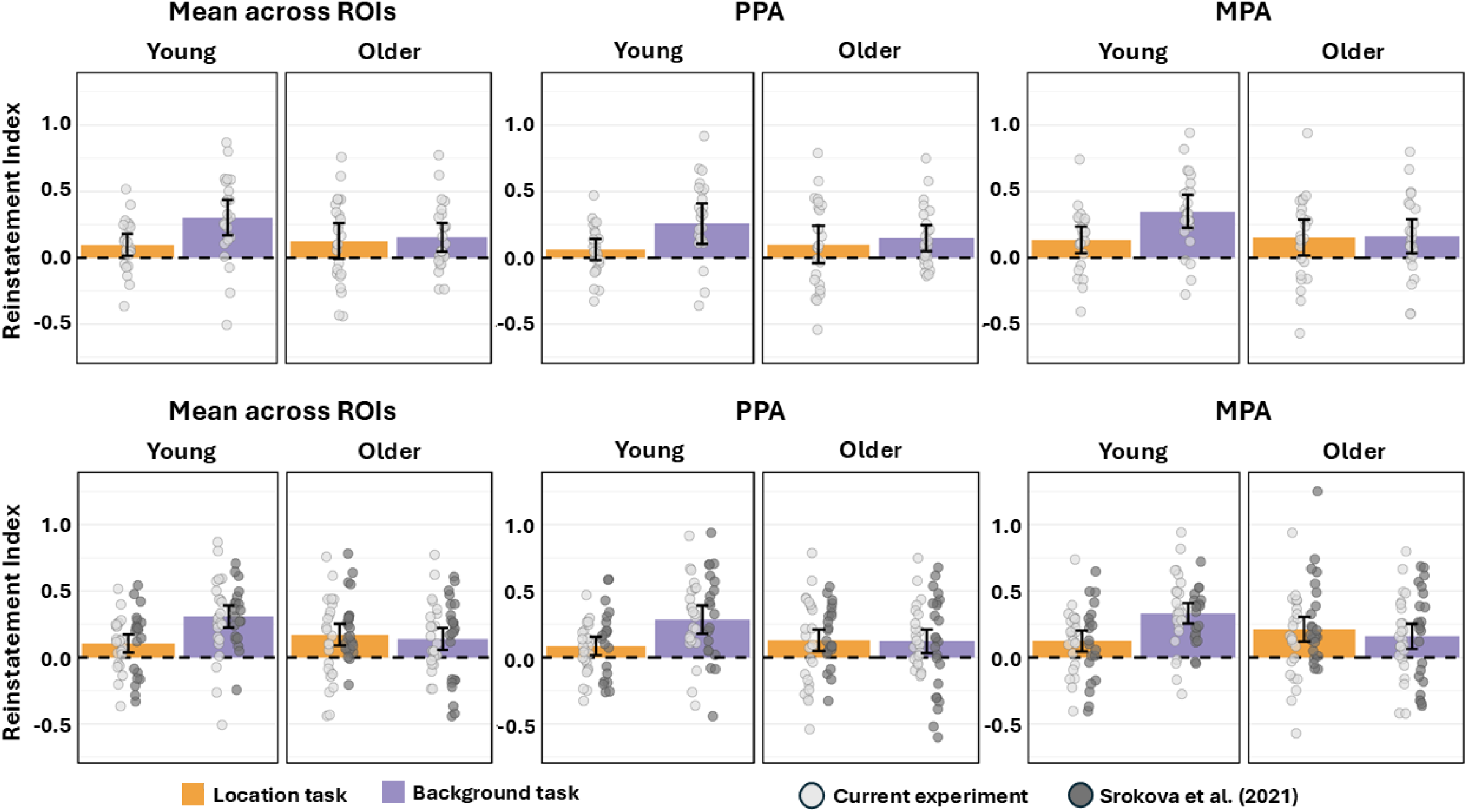
Plots of the reinstatement indices for items tested with the background and location tasks for the young and older groups for the current experiment (A) and for data pooled across experiments (B). Error bars represent 95% confidence intervals.

Given our pre-experimental hypothesis that, with a boost in memory performance for the background task, older adults would demonstrate reduced scene reinstatement in the location task relative to the background task (indicative of the ability to gate scene information when irrelevant to the task goal), we went on to examine reinstatement in young and older adults separately despite the non-significant age group by task interaction. Reinstatement effects were collapsed across hemispheres and ROIs, given the failure of these factors to moderate the initial findings. Paired samples t-tests revealed greater scene reinstatement in the background task relative to the location task in the young adults [t(23) = 2.784, p = 0.011, d = 0.568] but no such effect in the older age group [t(23) = 0.393, p = 0.698, d = 0.080] and, therefore, no evidence of retrieval gating in this group.

We went on to examine whether reinstatement effects (collapsed across hemispheres and ROIs) differed reliably from zero in each age group and task. Reliable effects were evident in both age groups for the location task [young: t(23) = 2.461, p = 0.011, d = 0.502; older: t(23) = 1.943, p = 0.032, d = 0.397] and the background task [young: t(23) = 4.714, p < 0.001, d = 0.962; older: t(23) = 3.006, p = 0.003, d = 0.614].

##### 3.3.1.1. Across experiments analyses

To directly compare the reinstatement indices derived from the current experiment with those from the experiment reported by Srokova et al. (2021), we ran a between experiments analysis (see Table 4, lower panel, for the data from the prior study), and Figure 4B for plots of the indices pooled across experiments). A 2 (experiment) x 2 (age group) x 2 (task) x 2 (ROI) x 2 (hemisphere) mixed effects ANOVA revealed a main effect of task [F(1,88) = 6.280, p = 0.014, partial η^2^ = 0.067], which was qualified by a significant age group by task interaction [F(1,88) = 11.529, p = 0.001, partial η^2^ = 0.116]. The three-way interaction between experiment, age group and task was not significant [F(1,88) = 0.670, p = 0.415, partial η^2^ = 0.008]. Following up on the age group by task interaction, pairwise t-tests (collapsed across experiments, ROIs and hemispheres) conducted separately for each age group indicated greater scene reinstatement in the background task relative to the location task in young adults [t(43) = 4.062, p < 0.001, d = 0.612] but no significant difference between the indices in the two tasks in older adults [t(47) = 0.653, p = 0.517, d = 0.094]. We also ran independent t-tests (again, collapsed across experiments, ROIs and hemisphere) to compare reinstatement indices between the two age groups for each task. The results revealed stronger reinstatement in the young relative to the older participants for the background task [t(90) = 2.865, p = 0.005, d = 0.598] but no significant difference between the indices for the location task [t(90) = 1.265, p = 0.209, d = 0.264]. In sum, these findings indicate that, when the data were pooled across the two experiments, scene reinstatement was robustly modulated by retrieval goal in the young but not the older adults and, additionally, that reinstatement in the background task was significantly attenuated in the older relative to the young age group.

To further elucidate these findings, we analyzed the magnitudes of the BOLD responses elicited by the test words studied with scenes vs. scrambled backgrounds (see Figure 5 for plots of the data pooled across experiments). We collapsed across ROIs and hemispheres as these factors did not interact with experiment, age group or task in the across-experiment analysis of the reinstatement indices. Thus, the data were subjected to a 2 (experiment) x 2 (age group) x 2 (background context: scenes, scrambled) x 2 (task) mixed effects ANOVA. The ANOVA revealed a main effect of background context [F(1,88) = 55.554, p < 0.001, partial η^2^ = 0.387], indicating greater activity in scene selective ROIs for words paired with scenes relative to scrambled backgrounds, a main effect of task [F(1,88) = 29.618, p < 0.001, partial η^2^ = 0.252], indicating greater activity in the background task relative to the location task, and a main effect of age group [F(1,88) = 14.433, p < 0.001, partial η^2^ = 0.141], indicating greater overall activity in older relative to young participants. There was also an interaction between experiment and task [F(1,88) = 25.296, p < 0.001, partial η^2^ = 0.223]. Follow up independent t-tests indicated that this interaction reflected greater overall activity for the previous experiment relative to the current experiment for the background task [t(90) = 3.535, p < 0.001, d = 0.738] but not for the location task (p = 0.977). The main effects were also qualified by interactions between context and task [F(1,88) = 6.702, p = 0.011, partial η^2^ = 0.071] and between age group, context and task [F(1,88) = 9.956, p = 0.002, partial η^2^ = 0.101]. As can be seen in Figure 5, the three-way interaction indicated significantly greater activity in older participants, relative to young adults, for items studied with scrambled backgrounds in both tasks (location task: [t(90) = 3.382, p < 0.001, d = 0.706]; background task [t(90) = 3.780, p < 0.001, d = 0.780]), as well as for scenes in the location task [t(90) = 4.089, p < 0.001, d = 0.853]. There were, however, no age differences in the background task for items that had been paired with scenes (p = 0.098). Additionally, context effects (scene > scrambled) were modulated by task in the young [F(1,42) = 12.905, p < 0.001, partial η^2^ = 0.235] but not the older participants (p = 0.648).

**Figure 5.**
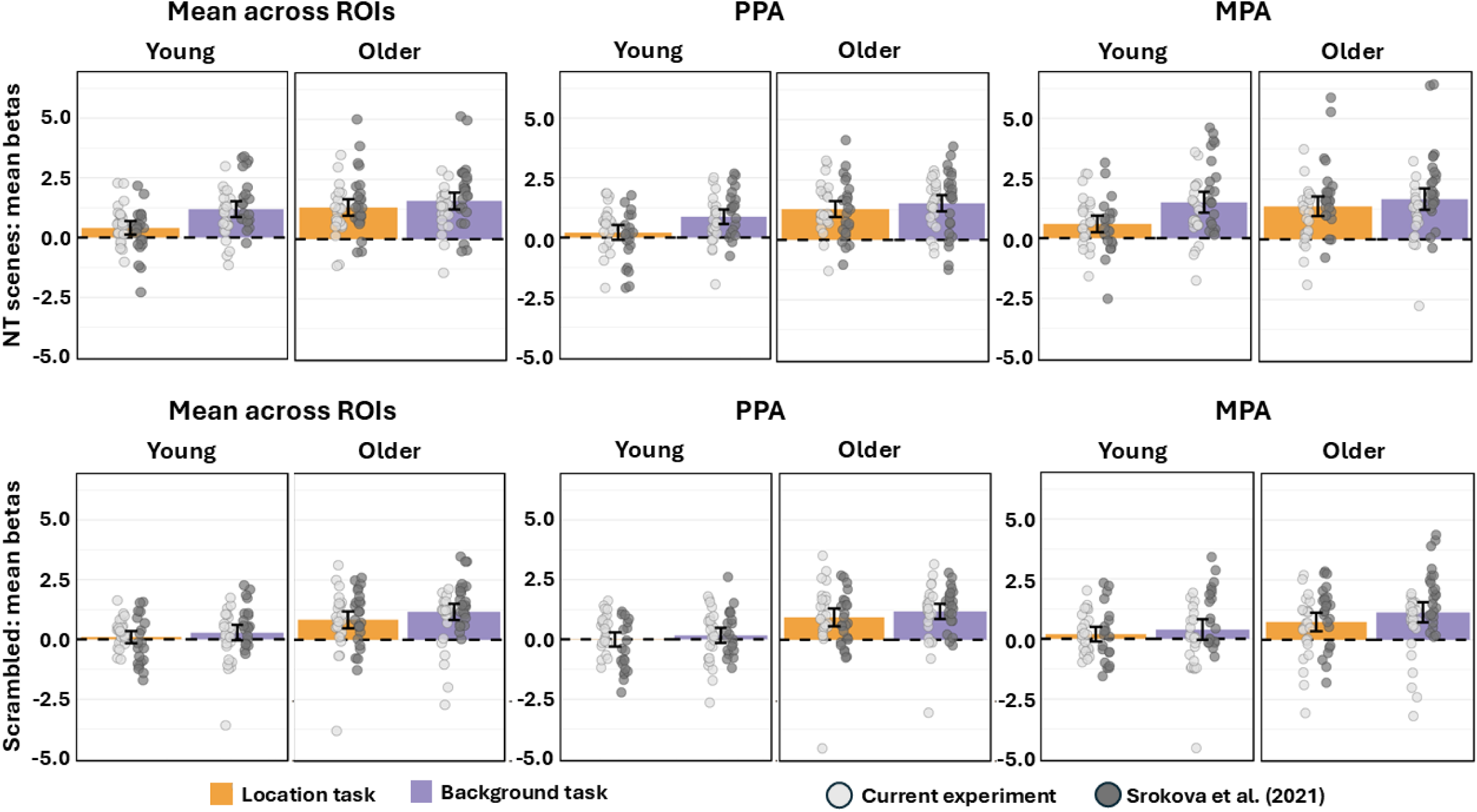
Plots of the beta estimates for items tested with the location and background tasks and studied with non-target scenes (A) and scrambled backgrounds (B) for the young and older groups pooled across experiments. Error bars represent 95% confidence intervals.

##### 3.3.1.2. Analysis restricted to source correct trials

For the reasons outlined in the Methods (section 2.9) the analyses of the effects of task and age group on scene reinstatement effects described above were conducted using all ‘non-target‘ test trials associated with test items correctly endorsed as studied. A key finding from these analyses was the age-related reduction in scene reinstatement evident in the background task. To rule out the possibility that this finding reflected age differences in the relative proportions of trials associated with correct vs. incorrect source memory judgments, we examined the effects of age on scene reinstatement in the background task with an analysis of the combined datasets that was restricted to trials associated with accurate judgments. We estimated reinstatement indices for all participants with 5 or more correct ‘scene’ and 5 or more correct ‘scrambled’ trials, leading to the loss of 1 older participant from the present experiment and 3 older participants from the prior study. The indices were subject to a 2 (experiment) x 2 (age group) x 2 (ROI) ANOVA. This revealed a single effect of age group (F1,84) = 6.076, p = 0.016, partial η^2^ = 0.067) that reflected lower reinstatement in the older age group (means (sds) collapsed across ROIs of 0.193 (0.361) and 0.380 (0.358) for older and younger groups respectively).

##### 3.3.1.3. Analyses after matching age groups for background reinstatement

The analyses reported in section 3.3.1.1. above revealed that, along with their failure to demonstrate retrieval gating, the older adults also demonstrated significantly weaker reinstatement than the young age group in the background task. This raises the possibility that the absent gating effects in the older group are merely a consequence of the fact that there was relatively little retrieved scene information available to be gated. To examine this possibility, we formed sub-samples of young and older adults that were matched for strength of scene reinstatement (collapsed across ROIs, hemispheres and experiment) in the background task (Ns = 38 per sub-group, mean (SD) reinstatement indices of 0.246 (0.239) and 0.248 (0.210) for the young and older subgroups respectively – see Figure 6). Paired t-tests revealed a robust gating effect in the young adults [t(37) = 2.966, p = 0.005, d = 0.481] but no evidence for an effect in the older sub-group [t(37) = 1.127, p = 0.267, d = 0.183]. Thus, the failure on the part of older adults to demonstrate retrieval gating cannot be attributed to relatively weak reinstatement in the background task.

**Figure 6.**
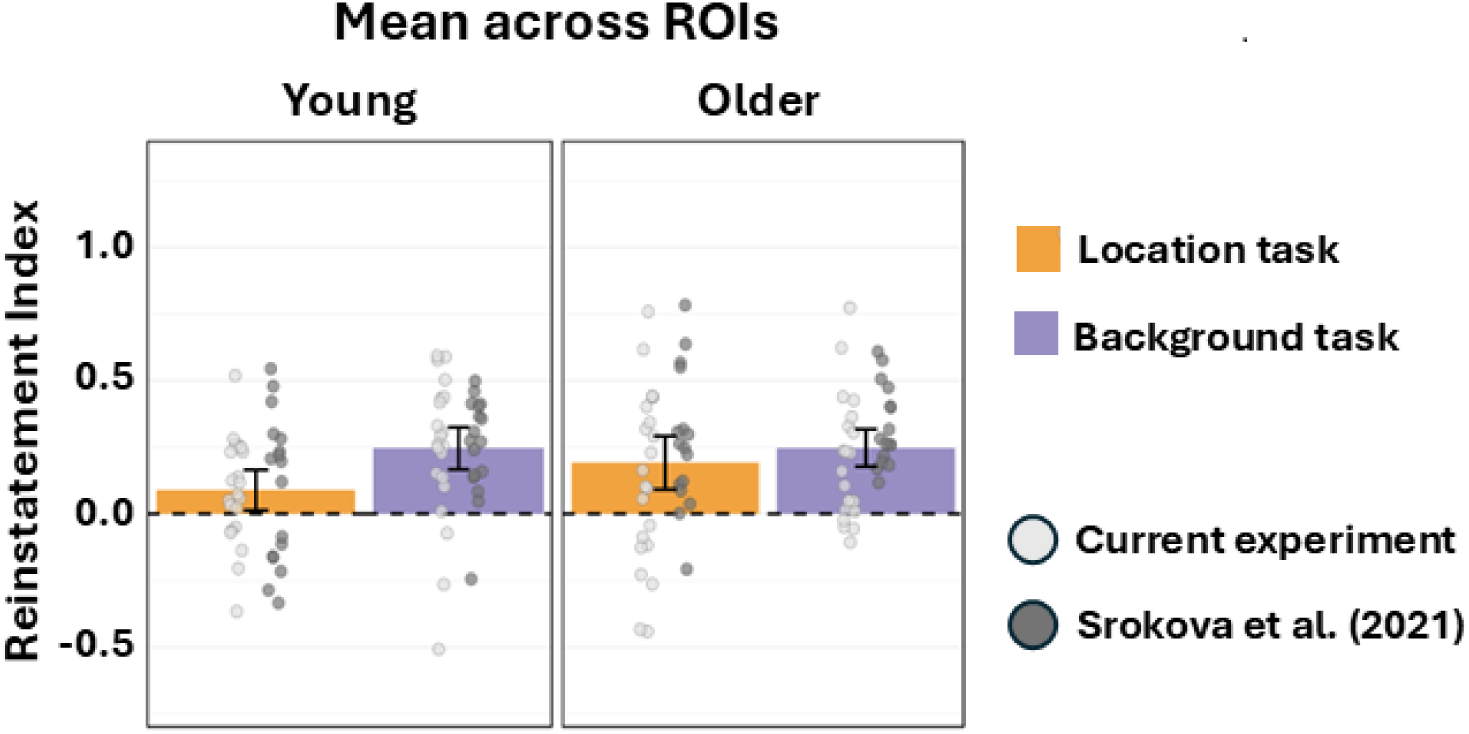
Plots of the reinstatement indices for items tested with the background and location tasks for subgroups of the young and older adults, matched for strength of scene reinstatement in the background task and collapsed across ROIs, hemisphere and experiment (ns = 38 per subgroup). Error bars represent 95% confidence intervals.

##### 3.3.1.4. Relationship between PPA scene reinstatement and memory performance

As was noted in the Introduction, Srokova et al (2021) reported positive associations across participants between PPA scene reinstatement in the background task and both source memory accuracy in the same task and item memory accuracy across tasks. Accordingly, we employed the same analysis approach here in an effort to replicate these findings (see Methods, section 2.10, for details of the regression model). Unlike in the prior study, here we were unable to identify any association between either item or source memory performance and PPA reinstatement or its interaction with age group (min p = 0.256). The same held true for the regression analysis predicting source memory from the combined datasets (see Methods; min p = 0.095). By contrast, in the combined model predicting item memory performance (Pr), the reinstatement by age group interaction term was reliable (β = -0.401, p = 0.002). Accordingly, follow-up analyses were conducted separately for each age group. That for the older group revealed no evidence for a predictive role of reinstatement (min p = 0.636). In the case of the young age group, however, the model revealed a significant reinstatement x experiment interaction term (β = 0.538, p = 0.003). Bivariate correlations conducted separately for each experiment reproduced the original finding of a highly robust association in the study of Srokova et al. (2021) but revealed a non-significant association with performance in the current experiment (r = 0.794, p < 0.001 and r = 0.273, p = 0.196, respectively).

## 4. Discussion

This study investigated the ability of young and older adults to employ retrieval gating to attenuate episodic content that is irrelevant to a retrieval goal. Specifically, we followed up a previous report that, unlike younger adults, older adults do not gate goal-irrelevant scene information, as indexed by their failure to modulate retrieval-related scene reinstatement (Srokova et al., 2021). Here, we sought to determine whether this failure was a consequence of the relatively weak memory for scene information demonstrated in that study by the older adults. Thus, we strengthened scene memory relative to the prior study by presenting study words and their associated background contexts twice during the encoding phase. Scene reinstatement was identified in the PPA and MPA in both age groups. However, despite a substantial boost to scene memory performance, it remained the case that only younger participants demonstrated retrieval gating effects. We elaborate on these and our other findings below.

### 4.1. Behavioral findings

Vividness ratings and RTs across the two study phases were statistically equivalent for the young and older groups. Therefore, any differences in the magnitude and pattern of scene reinstatement between the two age groups are unlikely to have been confounded by the vividness or speed with which the scenarios were imaged at study. In both age groups, vividness ratings were lower for words paired with scrambled backgrounds than for scenes and, in the first study phase, RTs were faster for items paired with scrambled backgrounds relative to those paired with scenes. These findings most likely reflect the fact that scenes contain richer information than scrambled backgrounds.

Unlike in Srokova et al. (2021), item memory in the present study was higher for the young than for the older participants. However, this difference was small and no longer significant when visual acuity was included as a covariate, adding to prior evidence that memory performance in older adults can be moderated by low-level perceptual factors (e.g., Davidson et al., 2018; Gellerson et al., 2023). In contrast, and consistent with the findings of Srokova et al. (2021) and numerous other reports (see Koen and Yonelinas, 2014, for review), source memory performance was substantially higher in the young than in the older adults.

The hypothesis motivating the present experiment was predicated on the success of the study manipulation in increasing older adults’ memory performance for background information above that reported in Srokova et al. (2021). The manipulation was successful in that source memory performance in the background task was robustly higher than in the prior study. Indeed, the memory performance of the older age group in the background task numerically (but non-significantly) eclipsed that of the young adults in Srokova et al. (2021). Thus, if the absent retrieval gating effects reported in that study for the older age group were indeed a consequence of age differences in memory for background information (see Introduction), gating would be expected to be evident in the older group in the present study.

### 4.2. fMRI

Turning first to the fMRI findings from the present experiment, it is clear that the experimental hypothesis was unsupported: there was a reliable gating effect in the young age group, but no sign of an effect in the older adults. Thus, the findings constitute a replication of those originally reported by Srokova et al. (2021) and negate the proposal that the prior results reflected a confound between age and background memory strength.

The across-experiments analysis of scene reinstatement revealed no evidence for a moderating effect of experiment on the effects of task or age group, or on the interaction between these factors. Accordingly, we take advantage of the additional statistical power gained by the near doubling of the sample sizes of the individual experiments and focus henceforth on the outcomes of the analysis of the combined datasets. These analyses identified a highly robust retrieval gating effect in the young participants, as expected given the results from the independent analyses, along with a null effect in the older age group. They also revealed a hitherto unappreciated aspect of the absent gating effects in the older adults. The lack of gating in these participants was associated, not with a failure to attenuate scene reinstatement in the location task, but with weaker reinstatement in the background task, as indicated by the combination of the robust age difference evident in reinstatement strength in the background task and the null age effect in the location task (see Figure 4). Of importance, the effect of age on reinstatement in the background task is not attributable to age differences in memory performance: the reinstatement effects evidenced by the young participants in the prior study were greater than those of the present older participants (t(42) = 2.206, p = 0.033, d = 0.668) despite the two groups demonstrating equivalent background memory performance. Equally relevant, a reliable age difference in background reinstatement remained when the analysis was restricted to trials associated with correct source judgments (see Results, section 3.3.1.2).

The finding of an age-related weakening of scene reinstatement in scene selective cortex is consistent with a previous report of age differences in retrieval-related scene reinstatement (Hill et al., 2021). In that study, participants were scanned both while they undertook a source memory test (discriminating between test words paired at study with scenes vs. faces) and during the preceding study phase, permitting an assessment of the selectivity with which faces and scenes were represented in their respective ‘preferred’ cortical regions as they were encoded (MPA and PPA in the case of scenes). Hill et al. (2021) reported that, as here, scene reinstatement effects (indexed by multi-voxel pattern similarity) were weaker in the older than the younger participants. Neural selectivity for scenes was also weaker, however, during the study phase, a finding consistent with numerous prior studies examining the effects of age on scene selectivity (e.g., Koen et al. 2019; see Koen and Rugg, 2019, for review). Crucially, Hill et al. (2021) reported that after controlling for scene selectivity at study age differences in scene reinstatement were no longer detectable. This finding was interpreted by the authors as indicating that age-related weakening of scene reinstatement is a consequence of having encoded relatively degraded (de-differentiated) scene representations. We conjecture that the present results can be explained similarly. That is, had we obtained fMRI data during the study phases of the two experiments described here, we would have found weaker scene selectivity in the older age group that mediated the age differences in scene reinstatement evident at retrieval.

Age-related weakening in neural selectivity in category-selective neural regions can result either from neural ‘broadening’ (heightened responses elicited by exemplars of ‘non-preferred’ categories) or neural ‘attenuation’ (reduced responses to exemplars belonging to a ‘preferred’ category). Both of these patterns, as well as their combination, have been reported previously when stimulus items were directly presented to participants (Chee et al., 2006; Park et al., 2012; Koen et al., 2019; Srokova et al., 2020), but we are unaware of any published study examining this issue when the category information is retrieved (reinstated) from memory rather than directly perceived (although see Hill et al., 2021 for an analogous analysis applied to multi-voxel pattern similarity analysis). The analysis of the BOLD responses separately elicited by the test items associated at study with scenes and scrambled images unequivocally indicates that the weaker reinstatement evidenced in the older age group was driven by neural broadening, that is, by enhanced activity for items that had been paired with scrambled images (see Figure 5). It will be of interest to examine in future research whether a similar pattern is evident at encoding. The present finding is somewhat analogous to reports that neurons in the sensory cortices of older non-human animals demonstrate broader tuning curves (reduced selectivity) in comparison to their younger counterparts, a finding that has been attributed to age-related weakening of GABAergic inhibition (for review, see Koen and Rugg, 2019).

How do the findings discussed above advance the understanding of age differences in retrieval gating? Most obviously, and as already noted, the findings indicate these differences do not reflect age differences in the strength (accuracy) of source memory. Additionally, the findings could be interpreted as casting doubt on the proposal that age-related reductions in gating effects reflect a failure to appropriately engage inhibitory control processes. This interpretation follows from the finding that, in the analyses conducted on the entire older sample, scene reinstatement in the background task in older participants was no greater than that for the location task in *either* age group. Hence the lack of evidence for gating in this age group could be argued to be a consequence of there being nothing much to gate. This possibility is decisively refuted, however, by the subsidiary analyses examining gating effects in sub-samples of the age groups in which the strength of scene reinstatement in the background task was equated (see section 3.3.1.3. above). Whereas gating effects remained highly robust in the young sub-group, they were far from significant in the older adults. Based on these findings, the proposal that age-related attenuation of retrieval gating effects is at least partially attributable to age differences in the efficacy of inhibitory control remains highly plausible (see Introduction and Amer et al., 2022). That being said, it will be of considerable interest to see the results of analogous studies that employ background contexts, such as faces, that elicit comparable levels of reinstatement in young and older age groups (cf. Hill et al., 2021).

The attenuated background reinstatement effects in the older participants raise the question of what sort of information supported their source memory judgments. It is commonly assumed that retrieval-related reinstatement supports the representation of episodic content, and hence that stronger reinstatement is associated with more content-rich memories (e.g., Thakral et al., 2015). The present findings appear to be inconsistent with this assumption: as noted above, the older adults demonstrated weaker background reinstatement than younger participants even when memory performance was closely matched (for a striking example of the reverse finding – equivalent reinstatement in the face of marked differences in source memory accuracy – see Elward et al., 2021). Moreover, there was no evidence in the older age group of an across-participant association between reinstatement strength and memory accuracy. Among the possible reasons for this seeming dissociation between background memory performance and reinstatement, two stand out (and are not mutually exclusive). The first possibility arises from the finding that, while older participants’ reinstatement effects in the background task were weak, they were nonetheless robustly greater than zero (p = 0.003). It is possible that this amount of reinstatement was sufficient to provide the episodic content necessary to support source memory judgments. By this argument, the larger reinstatement effects evident in the younger age group are indicative of the retrieval of more episodic information than was the case for the older adults, but much of this information was either non-diagnostic of source or redundant. The second possibility is that the older participants’ source memory judgments depended to a larger extent on relatively abstract, conceptual information than did the judgments of the younger adults. Evidence supporting this possibility comes from findings that, while episodic memories in older adults are relatively deficient in perceptual detail, they contain preserved or even enhanced conceptual (gist) information (e.g., Abadie et al., 2021; Koutstaal and Schacter, 1997). For example, older adults might have supported their source judgments to a greater extent than younger participants by remembering the scenarios they generated at study in response to each word-image pair. Relatedly, we cannot rule out the possibility that there were systematic age differences in the strategies employed to generate these scenarios that led to the encoding of less perceptually detailed memory representations in the older than the young participants.

As noted above, we were unable identify an association between older adults’ scene reinstatement in the background task and their memory performance, as was also the case in Srokova et al. (2021). In the one notable departure from the previous findings, however, in the present experiment we were also unable to identify reliable associations between scene reinstatement and memory performance in the younger age group. These null findings are particularly striking given the large effect sizes in the prior experiment. Moreover, the findings are also inconsistent with those of Elward et al. (2021) who, like Srokova et al. (2021), reported a positive association in cognitively healthy young adults between PPA (and MPA) scene reinstatement and source memory performance in a very similar background task. Further research will be required to determine whether the null findings in the present experiment reflect a type II error or a consequence of the employment of repeated study presentations.

### 4.3. Limitations

The present experiment shares one of the limitations discussed in Srokova et al. (2021), namely, the cross-sectional nature of the design. Thus, age differences in retrieval gating cannot be unambiguously attributed to the effects of aging rather than to a confounding variable such as a cohort effect or differential selection bias (Rugg, 2017). The experiment also shares another of the limitations identified by Srokova et al. (2021), namely a relatively small sample size that limits the power to distinguish between small and null effects. This limitation is mitigated, however, by the combined analyses of the present dataset and the one acquired by Srokova et al. (2021), which reinforce the independent findings of null gating effects in each study (indeed, a Bayesian analysis (JASP Team, 2024) of the combined data estimated that the likelihood of a true null was some 5 times greater (BF10 = 0.192) than the alternative).

### 4.4. Conclusions

The analyses described above confirm that, unlike younger adults, older adults do not modulate the strength of the cortical reinstatement of scene information according to its goal relevance, that is, they do not demonstrate retrieval gating of the information. The present findings further demonstrate that this age difference is not merely a consequence of age differences in memory strength for scenes, and additionally, that it is associated with relatively weak scene reinstatement even when the scene information is goal relevant. Crucially, however, retrieval gating effects are evident in young but not older adults even when the two age groups are sub-sampled to match the strength of scene reinstatement when the information is goal relevant. We conclude that the failure of older adults to gate goal-irrelevant episodic content is likely a reflection of age-related decline in top-down inhibitory control.

## Acknowledgements

We are grateful to all our participants and the Brain Health Imaging Center for their assistance with fMRI data acquisition.

## References

1. Abadie M, Gavard E, Guillaume F. Verbatim and gist memory in aging. Psychol Aging. 2021;36(8):891–901. 10.1037/pag0000635.

2. Abdulrahman H, Henson RN. Effect of trial-to-trial variability on optimal event-related fMRI design: Implications for Beta-series correlation and multi-voxel pattern analysis. Neuroimage 2016;125:756–766. 10.1016/j.neuroimage.2015.11.009.

3. Amer T, Wynn JS, Hasher L. Cluttered memory representations shape cognition in old age. Trends Cogn Sci. 2022;26(3):255–267. 10.1016/j.tics.2021.12.002.

4. Ashburner J, Friston KJ. Nonlinear spatial normalization using basis functions. Hum Brain Mapp. 1999;7:254--266. 10.1002/(SICI)1097-0193.

5. Bailey IL, Lovie-Kitchin JE. Visual acuity testing. From the laboratory to the clinic. Vision Res. 2013;90: 2– 9. 10.1016/j.visres.2013.05.004.

6. Benton AL. Differential behavioral effects in frontal lobe disease. Neuropsychologia 1968;6:53–60. 10.1016/0028-3932(68)90038-9.

7. Campbell KL, Lustig C, Hasher L. Aging and inhibition: Introduction to the special issue. Psychol Aging 2020;35(5):605–613. 10.1037/pag0000564.

8. Chadick JZ, Zanto TP, Gazzaley A. Structural and functional differences in medial prefrontal cortex underlie distractibility and suppression deficits in ageing. Nat Commun. 2014;5:4223. 10.1038/ncomms5223.

9. Chee MW, Goh JO, Venkatraman V, Tan JC, Gutchess A, Sutton B, Hebrank A, Leshikar E, Park D. Age-related changes in object processing and contextual binding revealed using fMR adaptation. J Cogn Neurosci. 2006;18(4):495–507. 10.1162/jocn.2006.18.4.495.

10. Cocosco CA, Kollokian V, Kwan RKS, Evans AC. Brainweb: online interface to a 3D MRI simulated brain database. Neuroimage 1997;5:425.

11. Davidson PSR, Vidjen P, Trincao-Batra S, Collin CA. Older Adults’ Lure Discrimination Difficulties on the Mnemonic Similarity Task Are Significantly Correlated With Their Visual Perception. J Gerontol B Psychol Sci Soc Sci. 2019;74(8):1298–1307. 10.1093/geronb/gby130.

12. de Chastelaine M, Mattson JT, Wang TH, Donley BE, Rugg MD. The neural correlates of recollection and retrieval monitoring: relationships with age and recollection performance. Neuroimage 2016;138:64–175. 10.1016/j.neuroimage.2016.04.071.

13. de Chastelaine M, Mattson JT, Wang TH, Donley BE, Rugg MD. Independent contributions of fMRI familiarity and novelty effects to recognition memory and their stability across the adult lifespan. Neuroimage 2017;156:340–351. 10.1016/j.neuroimage.2017.05.039.

14. de Chastelaine M, Wang TH, Minton B, Muftuler LT, Rugg MD. The effects of age, memory performance, and callosal integrity on the neural correlates of successful associative encoding. Cereb Cortex 2011;21(9):2166–2176. 10.1093/cercor/bhq294

15. Delis DC, Kramer JH, Kaplan E, Ober BA. California verbal learning test. 2^nd^ Ed. San Antonio, TX: The Psychological Corporation; 2000.

16. Duarte A, Henson RN, Graham KS. The effects of aging on the neural correlates of subjective and objective recollection. Cereb Cortex 2008;18(9):2169–2180. 10.1093/cercor/bhm243

17. Dulas MR, Duarte A. The effects of aging on material-independent and material-dependent neural correlates of contextual binding. Neuroimage 2011;57(3):1192–1204. 10.1016/j.neuroimage.2011.05.036.

18. Dulas MR, Duarte A. The effects of aging on material-independent and material-dependent neural correlates of source memory retrieval. Cereb Cortex 2012;22(1):37–50. 10.1093/cercor/bhr056.

19. Dulas MR, Duarte A. Aging affects the interaction between attentional control and source memory: an fMRI study. J Cogn Neurosci. 2014;26(12):2653–2669. 10.1162/jocn_a_00663.

20. Dulas MR, Duarte A. Age-related changes in overcoming proactive interference in associative memory: The role of PFC-mediated executive control processes at retrieval. Neuroimage 2016;132:116–128. 10.1016/j.neuroimage.2016.02.017.

21. Duverne S, Motamedinia S, Rugg MD. Effects of age on the neural correlates of retrieval cue processing are modulated by task demands. J Cogn Neurosci. 2009;21(1):1–17. 10.1162/jocn.2009.21001.

22. Elward RL, Rugg MD. Retrieval Goal Modulates Memory for Context. J Cogn Neurosci. 2015;27(12):2529–2540. 10.1162/jocn_a_00878.

23. Elward RL, Rugg MD, Vargha-Khadem F. When the brain, but not the person, remembers: Cortical reinstatement is modulated by retrieval goal in developmental amnesia. Neuropsychologia 2021;154:107788. 10.1016/j.neuropsychologia.2021.107788.

24. Fandakova Y, Shing YL, Lindenberger U. High-confidence memory errors in old age: the roles of monitoring and binding processes. Memory 2013;21(6):732–750. 10.1080/09658211.2012.756038.

25. Ferris FL, Kassoff A, Bresnick GH, Bailey IL. New Visual Acuity Charts for Clinical Research. Am J Ophthalmol. 1982;94(1):91–96. 10.1016/0002-9394(82)90197-0.

26. Gazzaley A, Clapp W, Kelley J, McEvoy K, Knight RT, D’Esposito M. Age-related top-down suppression deficit in the early stages of cortical visual memory processing. Proc Natl Acad Sci U S A. 2008;105(35):13122–13126. 10.1073/pnas.0806074105.

27. Gazzaley A, Cooney JW, Rissman J, D’Esposito M. Top-down suppression deficit underlies working memory impairment in normal aging. Nat Neurosci. 2005;8(10):1298–1300. 10.1038/nn1543.

28. Gellersen HM, McMaster J, Abdurahman A, Simons JS. Demands on perceptual and mnemonic fidelity are a key determinant of age-related cognitive decline throughout the lifespan. J Exp Psychol Gen. 2024;153(1):200–223. 10.1037/xge0001476.

29. Hill PF, King DR, Rugg MD. Age Differences In Retrieval-Related Reinstatement Reflect Age-Related Dedifferentiation At Encoding. Cereb Cortex 2021;31(1):106–122. 10.1093/cercor/bhaa210.

30. Horne ED, de Chastelaine M, Rugg MD. Neural correlates of post-retrieval monitoring in older adults are preserved under divided attention, but are decoupled from memory performance. Neurobiol Aging 2021;97:106–119. 10.1016/j.neurobiolaging.2020.10.010.

31. Jacoby LL, Shimizu Y, Velanova K, Rhodes MG. Age differences in depth of retrieval: Memory for foils. J Mem Lang. 2005;52(4):493–504. 10.1016/j.jml.2005.01.007.

32. Kim H. Dissociating the roles of the default-mode, dorsal, and ventral networks in episodic memory retrieval. Neuroimage 2010;50(4):1648–1657. 10.1016/j.neuroimage.2010.01.051.

33. King DR, de Chastelaine M, Elward RL, Wang TH, Rugg MD. Recollection-related increases in functional connectivity predict individual differences in memory accuracy. J Neurosci. 2015;35(4):1763–1772. 10.1523/JNEUROSCI.3219-14.2015.

34. Koen JD, Hauck N, Rugg MD. The Relationship between Age, Neural Differentiation, and Memory Performance. J Neurosci. 2019;39(1):149–162. 10.1523/JNEUROSCI.1498-18.2018.

35. Koen JD, Rugg MD. Neural Dedifferentiation in the Aging Brain. Trends Cogn Sci. 2019;23(7):547–559. 10.1016/j.tics.2019.04.012.

36. Koen JD, Yonelinas AP. The effects of healthy aging, amnestic mild cognitive impairment, and Alzheimer’s disease on recollection and familiarity: a meta-analytic review. Neuropsychol Rev. 2014;24(3):332–354. 10.1007/s11065-014-9266-5.

37. Koutstaal W, Schacter DL, Galluccio L, Stofer KA. Reducing gist-based false recognition in older adults: encoding and retrieval manipulations. Psychol Aging 1999;14(2):220–237. 10.1037//0882-7974.14.2.220.

38. Liu P, Hebrank AC, Rodrigue KM, Kennedy KM, Section J, Park DC, Lu H. Age-related differences in memory-encoding fMRI responses after accounting for decline in vascular reactivity. Neuroimage 2013;78:415–425. 10.1016/j.neuroimage.2013.04.053.

39. Maillet D, Rajah MN. Age-related differences in brain activity in the subsequent memory paradigm: a meta-analysis. Neurosci Biobehav Rev. 2014;45:246–57. 10.1016/j.neubiorev.2014.06.006.

40. Mattson JT, Wang TH, de Chastelaine M, Rugg MD. Effects of age on negative subsequent memory effects associated with the encoding of item and item-context information. Cereb Cortex. 2014;24(12):3322–3333. 10.1093/cercor/bht193.

41. Morcom AM, Rugg MD. Effects of age on retrieval cue processing as revealed by ERPs. Neuropsychologia 2004;42(11):1525–1542. 10.1016/j.neuropsychologia.2004.03.009.

42. Mumford JA, Turner BO, Ashby FG, Poldrack RA. Deconvolving BOLD activation in event-related designs for multivoxel pattern classification analyses. Neuroimage 2012;59(3):2636–2643. 10.1016/j.neuroimage.2011.08.076.

43. Nelson DL, McEvoy CL, Schreiber TA. The University of South Florida free association, rhyme, and word fragment norms. Beh Res Meth Ins C. 2004;36(3):402–407. 10.3758/BF03195588.

44. Park J, Carp J, Kennedy KM, et al. Neural broadening or neural attenuation? Investigating age-related dedifferentiation in the face network in a large lifespan sample. J Neurosci. 2012;32(6):2154–2158. 10.1523/JNEUROSCI.4494-11.2012.

45. Peirce J, Gray JR, Simpson, S, MacAskill, M, Höchenberger, R, Sogo, H, Kastman E, Lindeløv JK. PsychoPy2: Experiments in behavior made easy. Beh Res Methods 2019:51;195–203. 10.3758/s13428-018-01193-y.

46. Raven J, Raven JC, Courth JH. The advanced progressive matrices. In: Manual for Raven’s progressive matrices and vocabulary scales, Section 4. San Antonio: Harcourt Assessment; 2000.

47. R Core Team. R: a language and environment for statistical computing. Vienna: R Foundation; 2020.

48. Reitan RM, Wolfson D. The Halstead-Reitan neuropsychological test battery: therapy and clinical interpretation. Tucson: Neuropsychological; 1985.

49. Rugg MD. Retrieval processes in human memory: Electrophysiological and fMRI evidence. In: Gazzaniga MS, Ivry RB, Mangun GR, editors. The Cognitive Neurosciences, third ed. Cambridge MA: MIT Press; 2004. p. 727–738.

50. Rugg MD (2017). Interpreting age-related differences in memory-related neural activity. In: Cabeza R, Nyberg L, Park DC, editors. The cognitive neuroscience of aging: Linking cognitive and cerebral aging, 2nd ed. Oxford University Press; 2017. p. 183–203.

51. Rugg MD, Srokova S. Effects of age on neural reinstatement during memory retrieval. In: Grafman, JH, editor. Encyclopedia of the Human Brain, second ed, vol.4. USA: Elsevier; 2024. p. 189–201.

52. Rugg MD, Vilberg KL. Brain networks underlying episodic memory retrieval. Curr Opin Neurobiol. 2013;23(2):255–260. 10.1016/j.conb.2012.11.005.

53. Smith A. Symbol digit modalities test (SDMT) manual. Los Angeles: Western Psychological Services; 1982.

54. Snodgrass JG, Corwin J (1988). Pragmatics of Measuring Recognition Memory: Applications to Dementia and Amnesia. J Exp Psychol Gen. 1988;117(1):34–50.

55. Spreen O, Benton AL. Neurosensory center comprehensive examination for aphasia. Victoria: Neuropsychology Laboratory; 1977.

56. Srokova S, Hill PF, Elward RL, Rugg MD. Effects of age on goal-dependent modulation of episodic memory retrieval. Neurobiol Aging 2021;102:73–88. 10.1016/j.neurobiolaging.2021.02.004.

57. Srokova S, Hill PF, Koen JD, King DR, Rugg MD. Neural Differentiation is Moderated by Age in Scene-Selective, But Not Face-Selective, Cortical Regions. eNeuro. 2020;7(3):ENEURO.0142-20.2020. Published 2020 May 21. 10.1523/ENEURO.0142-20.2020.

58. Thakral PP, Wang TH, Rugg MD. Cortical reinstatement and the confidence and accuracy of source memory. Neuroimage 2015;109:118–129. 10.1016/j.neuroimage.2015.01.003

59. Wang WC, Cabeza R. Episodic memory encoding and retrieval in the aging brain. In: Cabeza R, Nyberg L, Park DC, editors. Cognitive Neuroscience of Aging: Linking Cognitive and Cerebral Aging, second ed. Oxford University Press, 2016. p. 301–336.

60. Wang TH, Johnson JD, de Chastelaine M, Donley BE, Rugg MD. The Effects of Age on the Neural Correlates of Recollection Success, Recollection-Related Cortical Reinstatement, and Post-Retrieval Monitoring. Cereb Cortex 2016;26(4):1698–1714. 10.1093/cercor/bhu333.

61. Wechsler D. WAIS-R: Wechsler adult intelligence scale-revised. New York: The Psychological Corporation; 1981.

62. Wechsler D. Wechsler test of adult reading. San Antonio: The Psychological Corporation; 2001.

63. Wechsler D. Wechsler memory scale, Ed 4. San Antonio: The Psychological Corporation; 2009.

64. Wechsler D. The test of premorbid function (TOPF). San Antonio, TX: The Psychological Corporation; 2011.

65. Weeks JC, Grady CL, Hasher L, Buchsbaum BR. Holding On to the Past: Older Adults Show Lingering Neural Activation of No-Longer-Relevant Items in Working Memory. J Cogn Neurosci. 2020;32(10):1946–1962. 10.1162/jocn_a_01596.

66. Wickham H. ggplot2: elegant graphics for data analysis. https://ggplot2-book.org/, 2016.

67. Yuan P, Raz N. Prefrontal cortex and executive functions in healthy adults: a meta-analysis of structural neuroimaging studies. Neurosci Biobehav Rev. 2014;42:180–192. 10.1016/j.neubiorev.2014.02.005.

68. Zanto TP, Gazzaley A. Aging of the frontal lobe. Handb Clin Neurol. 2019;163:369–389. 10.1016/B978-0-12-804281-6.00020-3.

